# Tunable Laminar Perfusion Coordinates Endothelial and Perivascular Remodeling in Angiogenic Vasculature-on-Chip

**DOI:** 10.64898/2026.07.21.739741

**Authors:** Tomasz Nawara, Katja Meier, Jan Kuom, Irene Hollfinger, Julia Kraxner, Katharina Koch, Maria Hastermann, Lenka Jablonicka, Laure Vinet Barancourt, Jennifer Schwarzkopf, Holger Gerhardt

**Affiliations:** Integrative Vascular Biology Laboratory, Max Delbrück Center for Molecular Medicine in the Helmholtz Association (MDC), Berlin 13125, Germany; German Center for Cardiovascular Research (DZHK), Berlin 10785, Germany; Experimental and Clinical Research Center (ECRC), a cooperation of Charité-Universitätsmedizin Berlin and Max Delbrück Center for Molecular Medicine, Berlin, Germany; Charité-Universitätsmedizin Berlin, corporate member of Freie Universität Berlin and Humboldt-Universität zu Berlin, Berlin, Germany; Berlin Institute of Health (BIH), Germany

**Keywords:** microfluidics, fluidic shear stress, micropump, mechanoadaptation

## Abstract

Perfusable vascular microphysiological systems are increasingly used to model angiogenesis, tissue crosstalk, and disease. However, many platforms still rely on oscillatory, discontinuous, or poorly controlled perfusion regimes, limiting the study of sustained flow-dependent vascular remodeling. Here, we establish a tunable, unidirectional laminar flow workflow for long-term perfusion of angiogenic vasculature-on-chip cultures and use it to investigate endothelial, perivascular, and immune cell responses to sustained flow. Using an AIM Biotech microfluidic platform containing 14-day-old human umbilical vein endothelial cell-derived angiogenic sprouts and pericytes, continuous perfusion enabled intraluminal transport of 1 μm tracer beads through vessels, demonstrating stable flow across the vascular bed. Sustained laminar flow induced endothelial remodeling at both the mother vessel and sprout levels, with cellular alignment evident in both compartments. Quantitative analysis of the mother vessel further revealed Golgi polarization against the direction of flow. Sustained perfusion also increased pericyte recruitment to angiogenic sprouts and reduced endothelial proliferation within the mother vessel, consistent with flow-driven vascular maturation and quiescence. Live-cell imaging further captured directional endothelial migration against the flow, lumen remodeling, and dynamic pericyte behavior under continuous perfusion. In immune-cell assays performed under continuous-flow conditions, interactions with untreated endothelium were limited, whereas inflammatory activation increased immune-cell adhesion and crawling. These observations suggest that sustained flow supports a quiescent endothelial phenotype and demonstrate the suitability of the platform for studying inflammatory activation and immune-vascular communication under controlled hemodynamic conditions. Beyond its biological relevance, the workflow provides practical advantages for live-cell imaging, low medium consumption, and downstream perturbation studies. Moreover, the modular design of the platform makes it well suited for vascular-organ crosstalk applications. Collectively, these results establish laminar flow angiogenic vasculature-on-chip as an experimentally tractable model for studying vascular mechanobiology, vascular maturation, and dynamic cell interactions under defined hemodynamic conditions.

## Introduction

Blood flow is a central regulator of vascular development, homeostasis, and disease^1, 2^. As blood passes through the vasculature, it generates fluid shear stress that varies in magnitude and profile between arteries, veins, and capillaries^3–5^. Endothelial cells, which line the inner surface of blood vessels, sense these mechanical cues and respond by remodeling their shape, polarity, migration, and transcriptional state^6–8^. Sustained hemodynamic forces also regulate vessel stability, lumen maintenance, network remodeling and regression, as well as interactions with mural cells^4, 9, 10^. However, when perturbed, fluid shear stress can also drive endothelial dysfunction and contribute to disease progression^3, 11^.

Much of our mechanistic understanding of these processes derives from 2D flow systems and animal-based experiments. While 2D systems have been crucial for dissecting core endothelial responses, they do not fully recapitulate the architecture, multicellular composition, and morphogenetic dynamics of angiogenic vessels^12–14^. Animal models remain essential for studying complex physiological responses and developing human therapeutics; however, they do not provide sufficient spatial and temporal resolution to capture subcellular dynamics in detail^15–17^. Furthermore, the ethical imperative and scientific need to reduce, refine, and ultimately replace animal use in biomedical research, as outlined by the 3R principles, have accelerated the development of alternative model systems^18–20^.

Microfluidic vasculature-on-chip systems provide a promising intermediate between reductionist monolayer assays and complex tissue models by enabling the formation of perfusable 3D vascular structures in controlled and imageable environments^21–23^. Within these systems, blood vessels can form through vasculogenesis, the *de novo* assembly of vessels from self-organizing endothelial cells, or through angiogenesis, the growth of vessels from preexisting vascular structures^24, 25^. In recent years, continuous perfusion and flow-driven remodeling have been demonstrated in several classes of vascular microphysiological systems, including preformed neovessel^26–28^ and prefabricated printed vessels models^29^, self-assembled microvascular networks^30, 31^, and vascularized tissue constructs^32^. However, less is known about how sustained laminar perfusion influences endothelial polarization, proliferation, lumen-associated remodeling, immune cell interactions, and mural cell recruitment within angiogenic sprout networks in experimentally tractable, low-volume systems compatible with routine live imaging^5^. This question is particularly important because angiogenic sprouts represent a highly dynamic vascular state in which lumenization, endothelial rearrangement, and perivascular stabilization occur simultaneously^9, 33^.

Here, we developed a modular perfusion workflow for imposing tunable, unidirectional, laminar flow across angiogenic vasculature-on-chip cultures and used it to test whether sustained flow is sufficient to drive mechanobiological adaptation and vascular maturation in established sprout networks. Using 14-day-old angiogenic sprout cultures composed of human umbilical vein endothelial cells (HUVECs) and supporting human brain pericytes, we show that continuous laminar perfusion supports intraluminal particle transport through sprout structures and induces canonical endothelial flow responses, including cellular alignment and Golgi polarization against the direction of perfusion. Sustained flow further enhanced pericyte recruitment to sprouts and reduced endothelial cell proliferation within the mother vessel. Through live-cell imaging, we additionally captured directional endothelial migration against the flow, a hallmark of flow-induced vascular remodeling^9^, dynamic pericyte migration, lumen formation, and immune cell crawling under perfusion conditions.

Together, our findings position laminar perfusion not merely as a maintenance condition for vascular chips, but as an instructive cue that reorganizes angiogenic vascular networks at both endothelial and multicellular levels. Importantly, the modular design of this workflow can be readily integrated into a broad range of microphysiological systems requiring controlled perfusion to establish physiological or pathological environments. These include vascularized organs-on-chip, functional blood-brain barrier models, and improved drug-delivery platforms.

## Results

### Tunable laminar flow supports sustained perfusion of angiogenic vasculature-on-chip

To develop a pump-based flow system capable of generating long-term, tunable, and laminar perfusion, we first established a microphysiological vascular network using the AIM Biotech idenTx 3 chip (**Fig. 1A**). In our design, Luer adapters enabled the use of standard Luer connectors, which are commonly employed in fluidic systems incorporating syringes or pumps^34, 35^. Because both the chips and adapters are commercially available, we selected this platform to benchmark the perfusion workflow presented in this study.

**Fig. 1.**
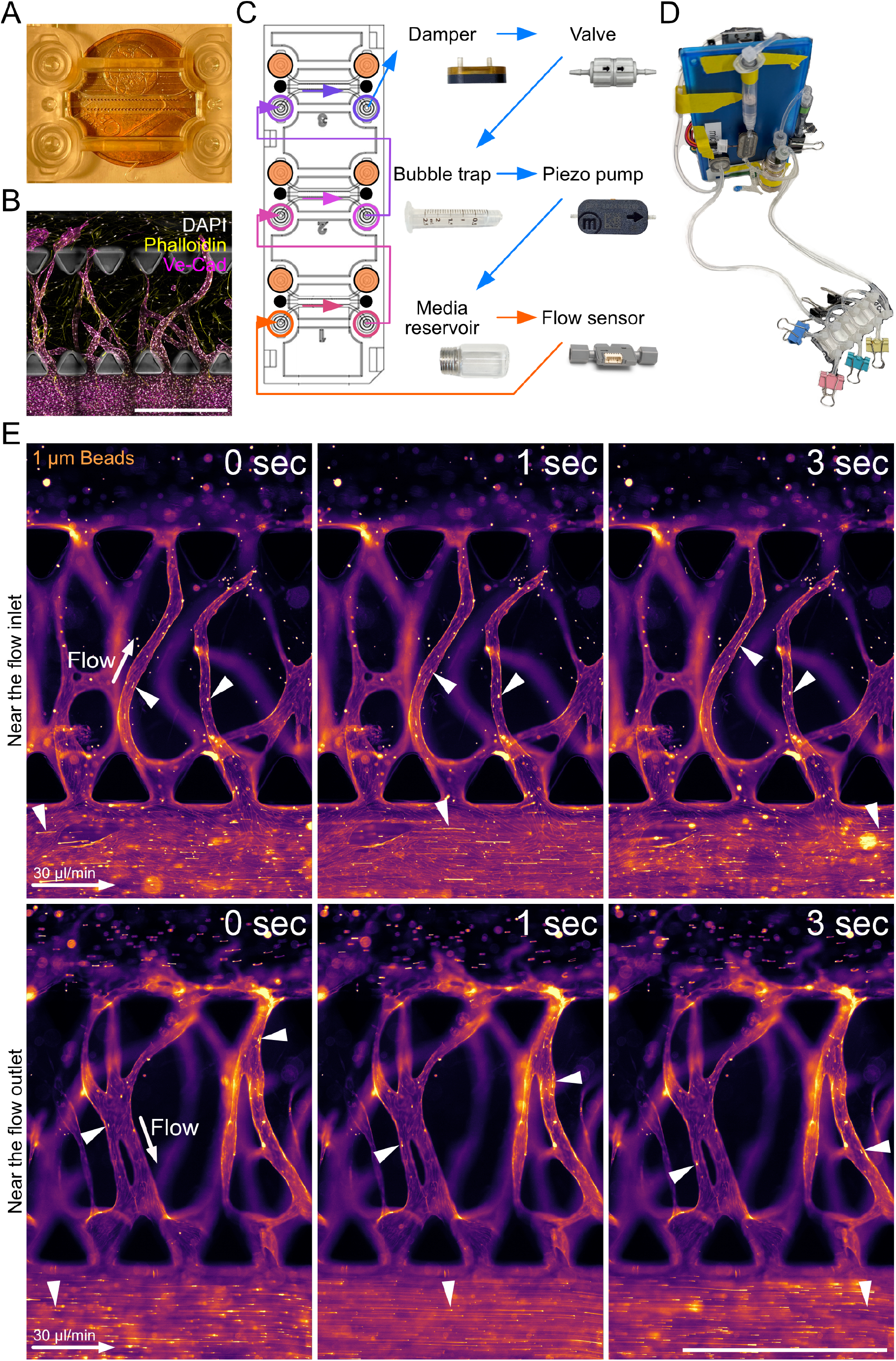
Tunable laminar-flow workflow enables sustained perfusion of angiogenic vasculature-on-chip. **A)** AIM Biotech idenTx 3 microfluidic chip. One of the three independent culture units is shown. A euro cent coin is included for scale. **B)** Spinning-disk confocal image of a 14-day-old angiogenic vasculature-on-chip culture established in the AIM Biotech platform. HUVECs were seeded into the bottom channel, a fibrin matrix was introduced into the central compartment, and human brain pericytes were seeded into the upper channel. Nuclei (DAPI, gray), F-actin (phalloidin, yellow), and Ve-Cadherin (Ve-Cad, magenta) are shown. Scale bar, 1 mm. **C)** Schematic representation of the closed-loop perfusion system. Medium passes sequentially through the three endothelial channels before entering a pulse damper, unidirectional valve, bubble trap, piezoelectric micropump, medium reservoir, and flow sensor before returning to the microfluidic chip. Gel-channel inlets were permanently sealed immediately after matrix polymerization (black circles), whereas pericyte-channel inlets were blocked at the adapter level prior to perfusion (orange circles). Colored arrows indicate the direction of medium flow. **D)** Photograph of the assembled perfusion platform showing the microfluidic chip, medium reservoir, flow sensor, pulse damper, unidirectional valve, bubble trap, and piezoelectric micropump. **E)** Representative time-lapse images showing transport of 1 μm fluorescent tracer beads through angiogenic sprouts during perfusion at 30 ìL/min. Arrowheads indicate bead trajectories within perfused vascular structures. HUVECs and tracer beads in orange. Scale bar, 1 mm.

The idenTx 3 chip is designed to accommodate three technical replicates. Each replicate consists of three pillar-separated channels. To establish a nascent angiogenic network, HUVECs were seeded into the left channel, serving as a proxy for the mother vessel, a thrombin-fibrinogen matrix was introduced into the central channel, and pericytes were seeded into the right channel. A nutrient gradient enabled HUVECs to invade the central matrix and collectively form lumenized sprouts over 14 days (**Fig. 1B**).

To investigate the effects of sustained flow on angiogenic vascular cultures, we developed a tunable, laminar, and unidirectional perfusion workflow compatible with microfluidic vasculature-on-chip systems. The platform was designed to support continuous flow, long-term culture compatibility, controllable flow rates, live-cell imaging access, and low total medium consumption, thereby enabling routine perfusion experiments in a standard benchtop setting. Continuous flow was achieved using a piezoelectric micropump. At the imposed flow rate of 30 µL/min, the Reynolds number in the mother vessel channel was estimated to be approximately 1.3–1.9, consistent with strongly laminar bulk flow. In this configuration, medium is pulled through the chip rather than pushed through it, minimizing the risk of leakage during long-term experiments.

Importantly, the gel-channel inlets were permanently sealed immediately after matrix polymerization on day 0, preventing fluid exchange through the gel-channel openings for the duration of the experiment. In contrast, the pericyte-channel inlets were blocked only at the adapter level immediately before perfusion. This prevented leakage and reservoir drying while maintaining medium flow through the pericyte channel during long-term experiments. In addition, the HUVEC channels were connected in series, enabling simultaneous perfusion of all three technical replicates within a single chip (**Fig. 1C**).

Starting from the microfluidic chip, the medium passes sequentially through the three HUVEC channels before entering a damper that reduces the intrinsic pulsatility generated by the micropump. The flow then passes through a unidirectional valve to prevent backflow and through a custom-made bubble trap before entering the piezoelectric micropump. After pumping, the medium flows through a reservoir and flow sensor before returning to the microfluidic chip (**Fig. 1D**). Because the system operates as a closed loop with approximately 5 mL of circulating medium, it requires substantially less medium than equivalent single-pass perfusion approaches, including conventional syringe-pump systems and recently developed waveform-generating platforms^28^.

We next asked whether established sprouts could be perfused under these conditions. Introduction of 1 μm tracer beads into the perfusion circuit revealed directional particle movement through the vascular sprout structures, indicating that the imposed flow was transmitted through the angiogenic network rather than being confined to the mother vessel (**Fig. 1E**, **Supplementary Movie 1.**, **Supplementary Movie 2.**). Bead transport was observed at a flow rate of 30 μL/min, demonstrating that the workflow supports sustained perfusion of the vascular bed while preserving optical accessibility for real-time imaging. Interestingly, sprouts located near the flow inlet experienced flow directed from the mother vessel toward the pericyte channel, whereas sprouts closer to the outlet exhibited reversed flow dynamics, with flow returning from the pericyte channel toward the mother vessel. Together, these data establish the technical basis for studying flow-dependent remodeling in angiogenic sprouts.

### Sustained laminar flow induces endothelial remodeling in angiogenic sprouts

We next asked whether endothelial cells within the vasculature-on-chip responded morphologically to sustained perfusion^6^. 14-day-old vascular cultures were maintained under static conditions or exposed to laminar flow at 30 μL/min for 24 h. Imaging of vascular endothelial-cadherin (Ve-Cad)-labeled endothelial cells revealed clear differences between static and flow-exposed cultures. Under flow conditions, endothelial cells within both the mother vessel and sprout compartments adopted a more elongated morphology, consistent with adaptation to a sustained directional mechanical cue (**Fig. 2A**). These observations suggest that endothelial cells embedded within angiogenic vascular networks remain mechanosensitive despite the geometric and multicellular complexity of the system. Thus, rather than serving solely as a means of medium exchange, laminar perfusion induced structural remodeling characteristic of endothelial flow adaptation within a three-dimensional organotypic environment.

**Fig. 2.**
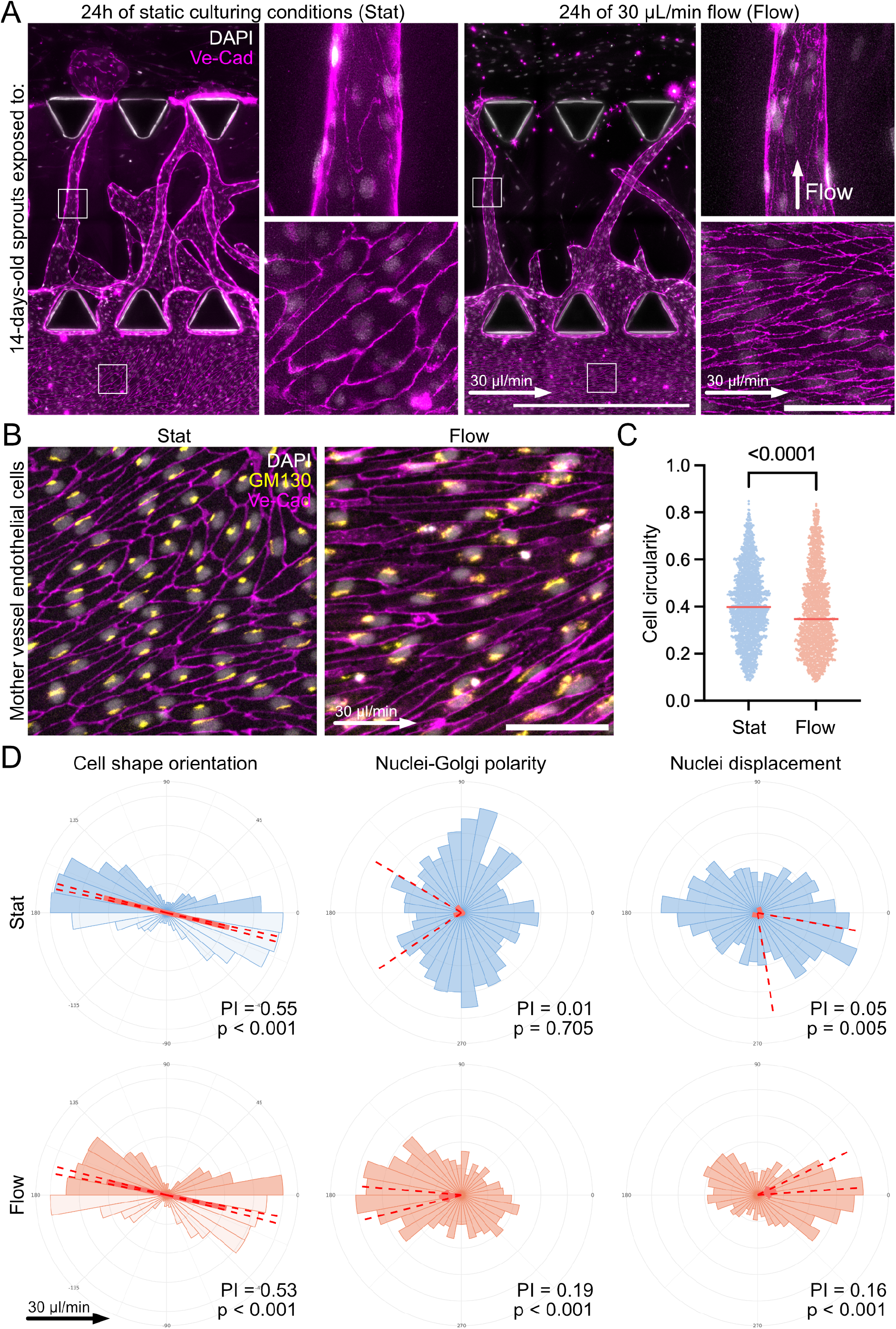
Sustained laminar flow induces endothelial remodeling and polarization. **A)** Representative immunofluorescence images of 14-day-old angiogenic vasculature-on-chip cultures maintained for an additional 24 h under static conditions (Stat) or exposed to laminar flow (Flow, 30 ìL/min). Insets highlight endothelial morphology within mother vessel and sprout regions. Perfusion direction is indicated by the white arrows. Nuclei (DAPI, gray) and Ve-Cadherin (Ve-Cad, magenta) are shown. Images were acquired using the MICA imaging platform (Leica Microsystems). Scale bars, 1 mm and 100 μm (insets). **B)** Representative images showing nuclei-Golgi polarity in HUVECs composing the mother vessel under static (Stat) and flow (Flow) conditions. Nuclei (DAPI, gray), Golgi (GM130, yellow), and Ve-Cadherin (Ve-Cad, magenta) are shown. Scale bar, 100 ìm. **C)** Quantification of HUVEC circularity under static (blue) and flow (orange) conditions. Data were not normally distributed according to the Shapiro-Wilk normality test. Red lines indicate median values (Stat = 0.397, Flow = 0.347). Circularity distributions differed significantly between conditions as determined by the Kolmogorov-Smirnov test (p < 0.0001). **D)** Rose plots showing endothelial alignment (left), nuclei-Golgi polarity (middle), and nuclear displacement (right) under static (blue) and flow (orange) conditions. Red arrows indicate the mean direction of the cell population, with arrow length corresponding to the polarity index. Red dashed lines indicate the 95% confidence intervals. The black arrow indicates the direction of flow relative to the flow dataset. Polarity indices and Rayleigh test statistics are shown. Rose plots were generated using the Polarity-JaM analysis pipeline. Quantitative analyses shown in (C-D) were obtained from three independent biological replicates comprising n = 2,010 cells under static conditions and n = 1,714 cells under flow conditions.

To quantify endothelial polarity responses under sustained perfusion, we analyzed cell circularity, cell orientation, nuclei-Golgi polarity, and nuclear displacement relative to the cell centroid within mother vessel regions using immunofluorescence labeling for Ve-Cadherin (Ve-Cad), golgin subfamily A member 2 (GM130), and DAPI (**Fig. 2B**), followed by Polarity-JaM-based analysis^36^. Under static conditions, endothelial cells displayed a more rounded morphology, reflected by increased cell circularity (**Fig. 2C**), while exhibiting a non-random distribution of cell orientations. In contrast, Golgi orientation remained largely unpolarized. Exposure to laminar flow resulted in a pronounced Golgi polarity bias against the direction of flow together with displacement of the nucleus toward the downstream side of the cell relative to the cell centroid (**Fig. 2D**). Across three biological replicates, the analysis included 2,010 cells under static conditions and 1,714 cells under flow conditions. The corresponding polarity indices and Rayleigh test statistics demonstrated a robust and reproducible endothelial polarization response to sustained laminar flow. Together, these findings show that sustained perfusion in this angiogenic vasculature-on-chip model is sufficient to induce a canonical endothelial mechanoadaptive program at the cellular level.

### Sustained flow enhances pericyte recruitment to angiogenic sprouts and reduces HUVECs proliferation

Because endothelial adaptation to flow is closely linked to vessel maturation, we next examined whether prolonged perfusion altered mural-cell association with the vascular network^9, 20^. Static and flow-exposed cultures were stained for Ve-Cad and platelet-derived growth factor receptor β (PDGFRβ) to visualize endothelial structures and associated pericytes. Flow exposure increased pericyte recruitment to angiogenic sprouts relative to static controls, as demonstrated by representative images (**Fig. 3A**) and quantification of pericyte number normalized to vessel area (**Fig. 3B**). This increase was consistently observed across three biological replicates and reached statistical significance. These findings identify enhanced pericyte recruitment as a multicellular consequence of sustained laminar perfusion and support the conclusion that flow promotes maturation of angiogenic vascular networks beyond its direct effects on endothelial morphology and polarity.

**Fig. 3.**
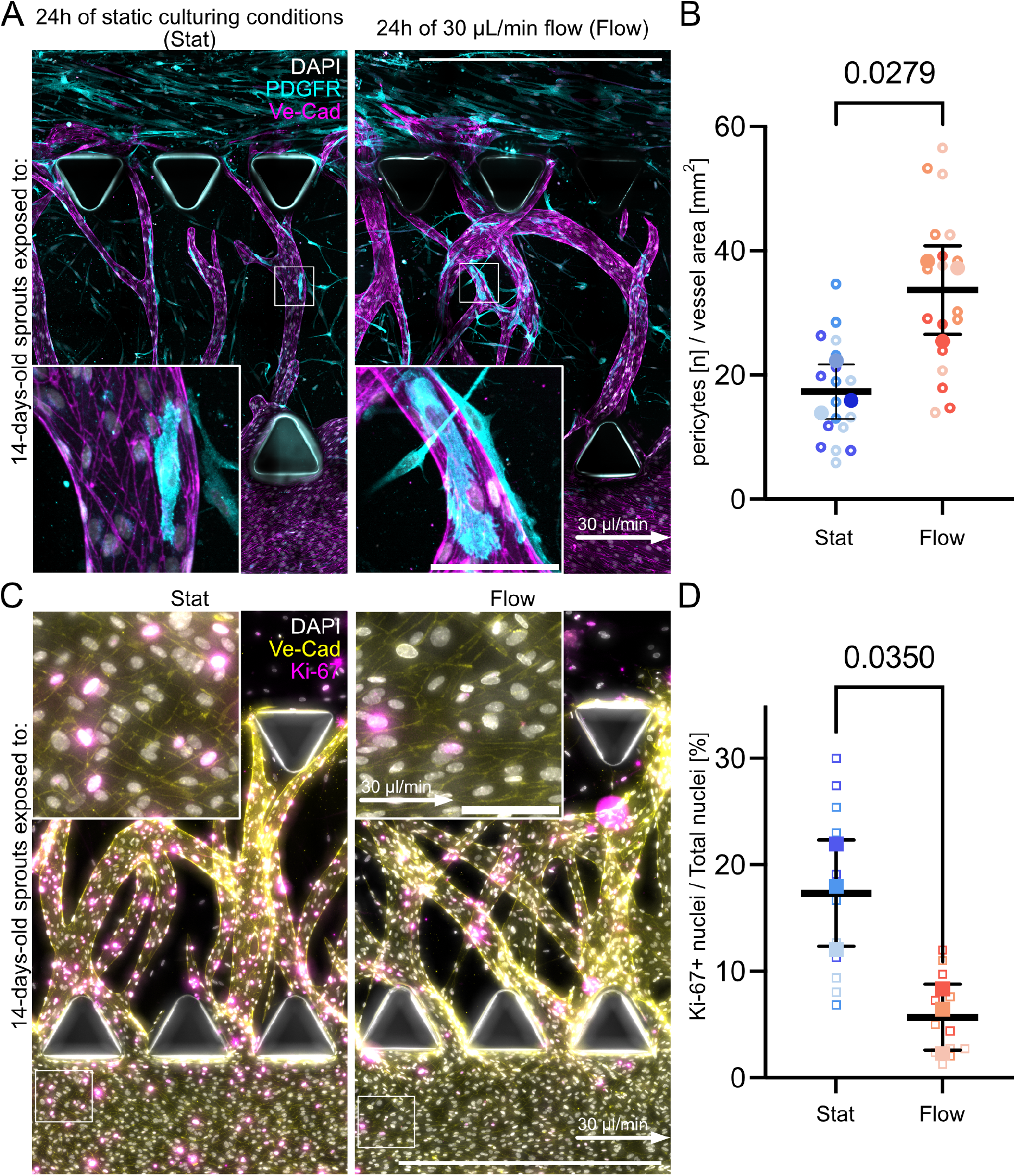
Sustained laminar flow promotes vascular maturation through increased pericyte recruitment and reduced endothelial proliferation. **A)** Representative immunofluorescence images of 14-day-old angiogenic vasculature-on-chip cultures maintained for an additional 24 h under static conditions (Stat) or exposed to laminar flow (Flow, 30 μL/min). Insets highlight HUVEC and human brain pericyte morphology within angiogenic sprouts. Nuclei (DAPI, gray), PDGFRβ-positive pericytes (cyan), and Ve-Cadherin (Ve-Cad, magenta) are shown. White arrows indicate the direction of perfusion. Images were acquired using spinning-disk confocal microscopy. Scale bars, 1 mm and 100 μm (insets). **B)** Quantification of vessel-associated pericytes normalized to vessel area. Each open circle represents one analyzed image, colors indicate independent biological replicates, and filled circles indicate the mean value for each biological replicate. Data were normally distributed according to the Shapiro-Wilk normality test and means per biological replicate were compared using a two-tailed unpaired t-test (p = 0.0279). Mean values were Stat = 17.33 and Flow = 33.66. Images used for quantification were acquired using the MICA imaging platform (Leica Microsystems). **C)** Representative images of Ki67 staining in static (Stat) and flow-exposed (Flow) cultures. Nuclei (DAPI, gray), Ve-Cadherin (Ve-Cad, yellow), and Ki67 (magenta) are shown. White arrows indicate the direction of perfusion. Images were acquired using the MICA imaging platform (Leica Microsystems). Scale bars, 1 mm and 100 μm (insets). **D)** Quantification of proliferating HUVECs within mother-vessel regions under static (Stat) and flow (Flow) conditions. Each open square represents one analyzed region of interest (ROI), colors indicate three technical replicates corresponding to the three culture units within a single microfluidic chip, and filled squares indicate the mean value for each technical replicate. Data were normally distributed according to the Shapiro–Wilk test, and technical-replicate means were compared using Welch’s t-test (p = 0.035). Mean values were Stat = 17.3% and Flow = 5.7%. Data were obtained from one biological experiment with three technical replicates per condition.

To assess whether sustained perfusion also affected the proliferative state of the vascular cultures, we quantified expression of the proliferation marker Ki67 within mother vessel endothelial cells (**Fig. 3C**). Flow exposure resulted in a reduction in endothelial proliferation, with the mean fraction of Ki67-positive HUVECs decreasing from 17.3% under static conditions to 5.7% following laminar perfusion (**Fig. 3D**). This reduction was observed across all analyzed culture units and reached statistical significance when technical-replicate means were compared (p = 0.035). Because these data were obtained from a single biological experiment, the result should be considered exploratory. Together with the increased pericyte recruitment, these findings are consistent with a shift from a highly proliferative angiogenic state toward a more mature and quiescent vascular phenotype under sustained flow.

### Live-cell imaging captures endothelial migration and lumen remodeling under flow

Our flow pipeline was designed to support live imaging experiments in perfused vasculature-on-chip systems. This is particularly important for studying subcellular protein dynamics, which remain difficult to capture in mammalian animal models because intravital microscopy does not provide sufficient spatial and temporal resolution^37, 38^. To determine whether sustained flow continues to support angiogenic activity, we performed live-cell imaging of vasculature-on-chip cultures exposed to laminar perfusion (**Fig. 4A**, **Supplementary Movie 3.**). Using label-free imaging, we captured HUVECs migration against the direction of flow both within the mother vessel and in angiogenic sprouts (**Fig. 4B**, **Supplementary Movie 4.**, **Supplementary Movie 5.**, **Supplementary Movie 6.**). In addition, we observed several dynamic multicellular behaviors, including lumen formation and tip-cell extension under flow conditions, as well as pericyte projection extension toward the sprout (**Fig. 4C**, **Supplementary Movie 7.**). Together, these observations extend the role of laminar perfusion beyond a purely morphological stimulus and suggest that sustained flow actively influences vascular maintenance, remodeling, and maturation across multiple biological scales.

**Fig. 4.**
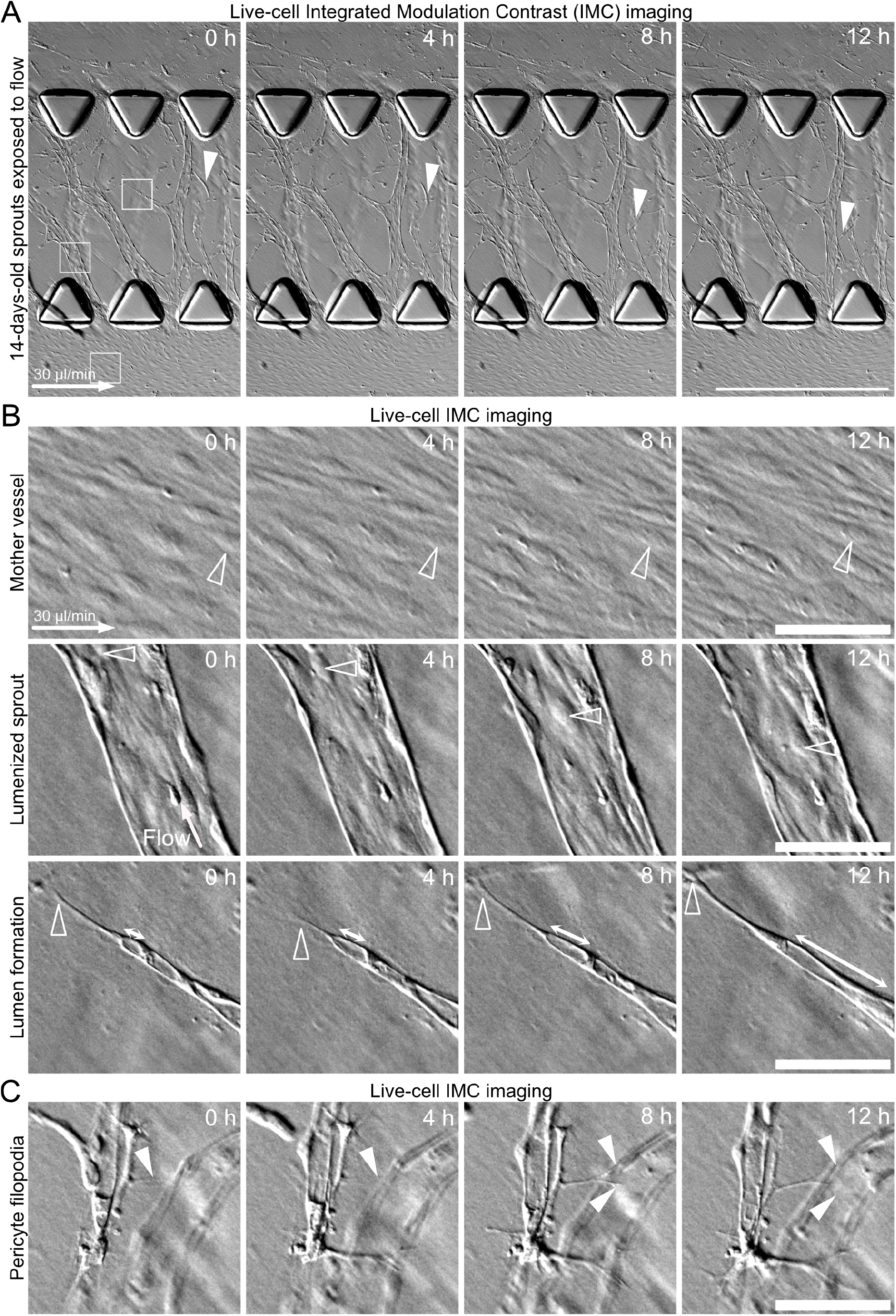
Live-cell imaging reveals dynamic endothelial and perivascular remodeling under flow. **A)** Representative time-lapse images acquired using integrated modular contrast (IMC) showing endothelial and pericyte dynamics within perfused 14-day-old vasculature-on-chip cultures. White filled arrowheads indicate migrating pericytes. White squares indicate regions enlarged in (B). White arrows indicate the direction of perfusion. Scale bar, 1 mm. **B)** Enlarged views of dynamic endothelial behaviors observed under sustained laminar flow. White open arrowheads indicate HUVEC migration against the direction of flow within the mother vessel (top) and angiogenic sprout (middle), as well as lumen formation within the sprout compartment (bottom). Scale bar, 100 μm. **C)** Representative time-lapse sequence showing extension of pericyte projections toward angiogenic sprouts. White filled arrowheads indicate extending pericyte filopodia. Scale bar, 100 ìm.

### The perfusion workflow supports immune cell interaction assays under flow

The piezoelectric pump used in this study is compatible with the perfusion of particles smaller than 50 μm. To test whether the workflow could also support the circulation of suspended cells, peripheral blood mononuclear cells (PBMCs, 5 × 10^6^ cells/mL medium) were introduced into the medium reservoir. Before injection, PBMCs were stimulated with staphylococcal enterotoxin B (SEB^39^, 0.3 µg/mL), while the vasculature-on-chip cultures were stimulated for 24 h with TNFα (3.3 ng/mL). In addition, the perfusion medium contained TNFα (3.3 ng/mL) throughout the experiment^40^. Compared with unstimulated PBMCs perfused through untreated angiogenic sprouts in the absence of TNFα, stimulated PBMCs displayed increased crawling behavior within the mother vessel compartment. In contrast, PBMCs in non-treated control conditions only rarely associated with the endothelial structures (**Fig. 5A**, **Supplementary Movie 8.**, **Supplementary Movie 9.**). Importantly, crawling of PBMCs within angiogenic sprouts could also be observed under stimulated conditions (**Fig. 5B**, **Supplementary Movie 10.**). These initial observations of immune cell rolling and crawling under flow conditions indicate that the system can be used to study inflammatory and immune-vascular interactions within angiogenic sprouts and endothelial monolayers. The incorporation of immune cell assays substantially broadens the translational utility of the platform, demonstrating that the workflow supports not only studies of vascular maturation, but also dynamic multicellular interaction assays under defined hemodynamic conditions.

**Fig. 5.**
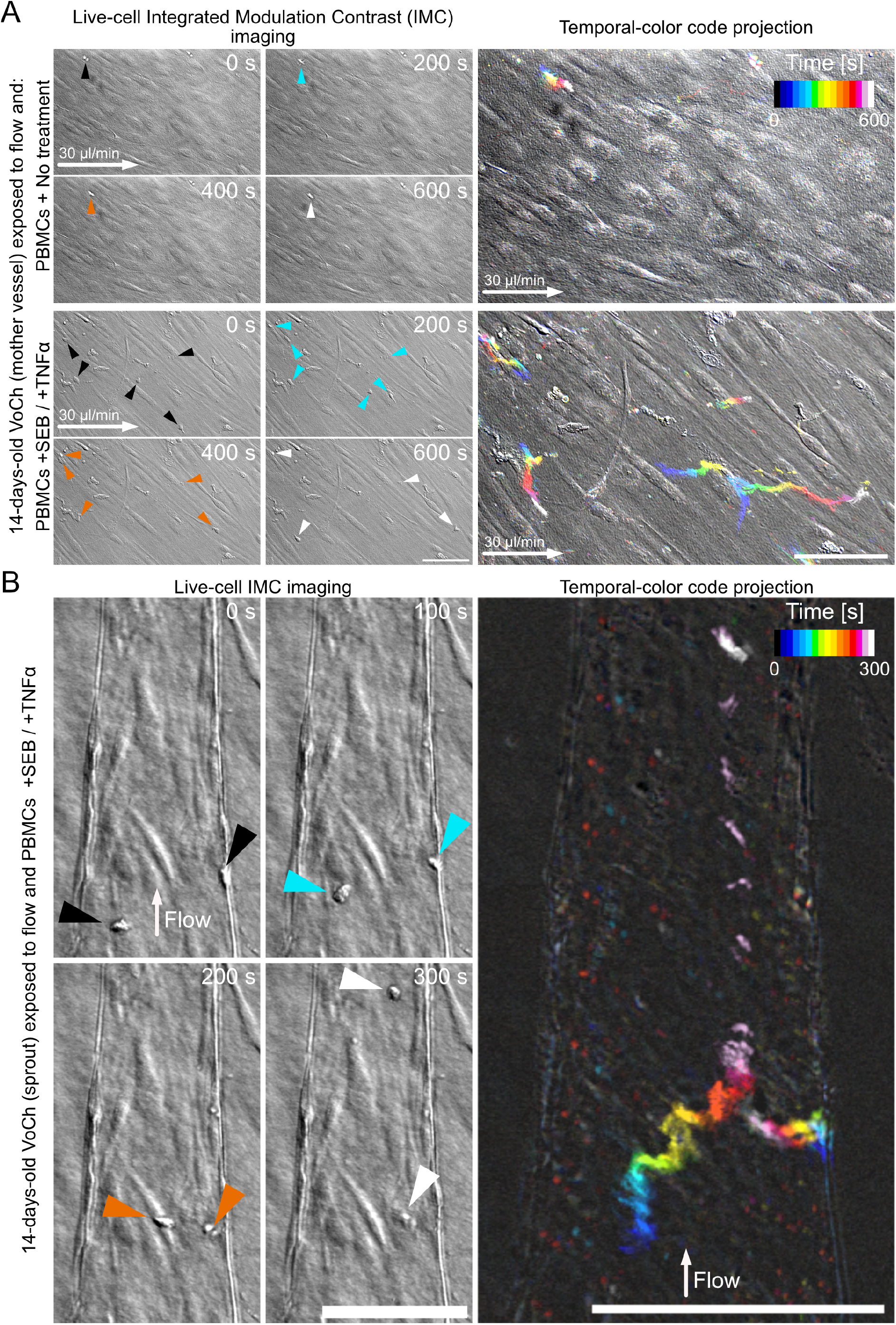
The perfusion workflow supports immune cell interaction assays under flow. **A)** Representative time-lapse images showing interactions between peripheral blood mononuclear cells (PBMCs) and endothelial cells within the mother vessel compartment under control (untreated; top) and inflammatory (bottom) conditions. White arrows indicate the direction of perfusion. PBMC crawling behavior throughout the imaging period is visualized using temporal color coding (right), where PBMCs present at the beginning of the imaging sequence are shown in blue, cells appearing at intermediate time points are shown in yellow, and cells present at later time points are shown in white. Corresponding colored arrowheads indicate representative PBMC trajectories within the time-lapse images. Under inflammatory conditions, PBMCs display increased endothelial association and crawling behavior compared with untreated controls. Scale bars, 100 μm. **B)** Representative images showing PBMC crawling under flow within an angiogenic sprout under inflammatory conditions. Arrowheads indicate PBMCs undergoing endothelial interactions and crawling events within the sprout compartment. White arrows indicate the direction of perfusion. Together, these observations demonstrate that the perfusion workflow supports immune-cell trafficking assays within both mother vessel and angiogenic sprout compartments under defined hemodynamic conditions. Scale bars, 100 μm.

## Discussion

In this study, we developed a tunable laminar-flow workflow for long-term perfusion of angiogenic vasculature-on-chip and demonstrate that sustained unidirectional flow acts as an instructive cue for vascular remodeling and maturation. Using an angiogenic sprout model based on the commercially available AIM Biotech idenTx 3 platform, we show that continuous perfusion supports stable intraluminal flow through developing vascular sprouts (**Fig. 1**) and induces coordinated endothelial and multicellular responses associated with vascular adaptation. These include endothelial elongation, alignment, Golgi polarization against the direction of flow (**Fig. 2**), increased pericyte recruitment, reduced endothelial proliferation (**Fig. 3**), lumen-associated remodeling (**Fig. 4**), and compatibility with immune-cell interaction assays (**Fig. 5**). Together, these findings position sustained laminar perfusion as a biologically active regulator of angiogenic vascular organization.

A central conceptual contribution of this work (**Fig. 1**) is therefore not simply that flow can be imposed on a vascular microphysiological system, as continuous perfusion has already been demonstrated in several vascular-chip platforms^5^, but rather that sustained laminar flow is sufficient to drive canonical mechanoadaptive responses in an angiogenic sprout context. This distinction is important because angiogenic sprouts differ fundamentally from preformed endothelial tubes and stabilized self-assembled vascular networks^41^. Sprouting vessels represent a highly dynamic developmental state characterized by endothelial rearrangement, active migration, lumen formation, proliferative activity, and ongoing interactions with mural cells and extracellular matrix^9^. Our findings indicate that sustained laminar flow can organize this plastic vascular state toward a more mature phenotype while preserving accessibility for live imaging and experimental perturbation.

The endothelial polarity analyses (**Fig. 2**) provide mechanistic links between this platform and established principles of vascular mechanobiology. Golgi polarization against the direction of flow is a hallmark of endothelial shear-stress sensing and coordinated migratory adaptation^6, 7, 9^. While this response has been extensively characterized in two-dimensional flow systems and *in vivo* vascular beds, demonstrating it within angiogenic vasculature-on-chip extends a canonical endothelial mechanoadaptive readout into a three-dimensional organotypic setting. The observation that sprouts near the inlet and outlet experienced opposite flow directions further suggests that angiogenic networks generate spatially complex perfusion environments even within relatively simple microfluidic geometries. Such local flow heterogeneity may become biologically relevant for future studies investigating tip-cell dynamics, lumen remodeling, or endothelial competition under flow^5^.

Beyond endothelial polarity, sustained perfusion also promoted multicellular features associated with vascular maturation (**Fig. 3**). Increased pericyte recruitment to angiogenic sprouts suggests that flow supports stabilization of endothelial-perivascular interactions, consistent with observations from developmental and physiological vascular remodeling *in vivo*^42–44^. Similarly, the strong reduction in HUVEC proliferation within the mother vessel compartment under flow conditions is consistent with the transition from a highly proliferative angiogenic state toward a more quiescent and mature endothelial phenotype^29, 45^. Together, these observations support a model in which sustained laminar perfusion coordinates endothelial and mural-cell adaptation across multiple biological scales.

The live-imaging compatibility of the platform further expands its utility (**Fig. 4**). Using label-free imaging, we were able to capture endothelial migration against the direction of flow, dynamic pericyte behavior, lumen formation, and tip-cell extension within living angiogenic sprouts. Such subcellular and multicellular dynamics remain difficult to resolve in mammalian animal models because intravital microscopy often lacks the spatial and temporal resolution required to monitor fine structural remodeling over extended periods^37, 38^. Thus, the present workflow provides an experimentally tractable intermediate between reductionist *in vitro* systems and complex *in vivo* models while remaining compatible with the ethical and scientific goals of reducing animal experimentation.

The immune cell perfusion experiments further demonstrate the versatility of the system (**Fig. 5**). Stimulated PBMCs exhibited increased crawling and endothelial association under inflammatory flow conditions. Although these datasets remain exploratory, they indicate that the platform can support studies of immune-vascular interactions under defined hemodynamic conditions. This substantially broadens the translational relevance of the workflow and suggests future applications in inflammation, vascular immunology, cancer biology, and therapeutic delivery studies.

Recent advances in vascular microphysiological systems have significantly improved the ability to establish long-term perfusion and flow-driven remodeling *in vitro*. Continuous physiological flow has been shown to rescue regressing self-assembled vascular networks, prolong vascular stability^31^, and support computationally informed analyses of hemodynamics and endothelial transcriptional adaptation^32^. Other studies established perfusable neovessel systems embedded within fibrin matrices and demonstrated their utility for barrier-function assays and immune-cell perfusion experiments^26^. More recently, advanced vascularized tissue platforms such as VIVOS extended intraluminal perfusion to millimeter-scale vascular beds and engineered tissues under physiological pressure and shear-stress conditions^32^. In parallel, the Hemadyne platform has enabled rapid reproduction of complex physiological and pathological flow waveforms and sustained endothelial culture for up to 60 days^28^. Together, these studies established sustained flow as a critical regulator of engineered vascular systems.

Our platform differs from these systems primarily in biological focus and experimental accessibility. Whereas many previous models emphasized stabilization of preformed vessels, rescue of self-assembled microvascular networks, or large-scale tissue vascularization, our workflow specifically interrogates the angiogenic sprout state. This focus enables simultaneous investigation of endothelial polarization, proliferation, lumen dynamics, mural-cell recruitment, and immune-cell interactions within the same experimentally tractable microfluidic format. Importantly, the workflow combines tunable long-term perfusion with low media consumption, live imaging compatibility, and modular assembly. In our setup, as little as 5 mL of media is sufficient for long-term operation, whereas conventional syringe-pump systems operating at 30 μL/min would require substantially larger media volumes over equivalent experimental durations. Such reductions are particularly important for studies involving expensive therapeutics, primary and patient-derived cells, cytokines, or long-term perturbation experiments.

The platform also incorporates several practical design considerations that improve experimental robustness and reproducibility. Within each idenTx 3 chip, all three HUVEC channels are connected in series, allowing simultaneous perfusion of all technical replicates under identical flow conditions. This is particularly important because substantial variability can occur between individual microfluidic channels, making it advantageous to use a single chip per experimental condition. In addition, the workflow was designed to support cleaning and recycling of major flow components, which enables straightforward reuse of the perfusion system between experiments. Most importantly, even though the platform was applied to AIM Biotech idenTx 3 chip, it can be easily adapted to virtually any microfluidic chip. Although the present study focused on a single flow regime (30 μL/min), the workflow is readily adaptable to a broad range of flow rates, enabling future studies investigating the effects of distinct hemodynamic conditions on vascular remodeling and disease.

Several limitations of the present study should be acknowledged. First, although the observed biological responses are consistent with sustained endothelial mechanoadaptation, we do not provide a detailed computational or experimental map of local shear stress across the vascular network. Local shear stress is expected to vary with sprout geometry, lumen dimensions, and network conductance. Future integration of particle-tracking velocimetry and computational fluid dynamics would enable more precise characterization of flow distribution and shear-stress profiles within individual angiogenic sprouts. Second, the reduction in endothelial proliferation was observed in a single biological experiment with three technical replicates and therefore requires confirmation across additional independent experiments. Third, while increased pericyte recruitment, reduced proliferation, and lumen-associated remodeling are all consistent with enhanced vascular maturation, additional functional readouts such as dextran permeability, barrier integrity measurements, basement membrane deposition, or direct lumen continuity analyses would further support this conclusion. Fourth, the current platform is not highly scalable and is primarily optimized for mechanistic and imaging-based experimentation rather than high-throughput screening applications. Finally, while our imaging-based analyses reveal coordinated changes in endothelial polarity, proliferation, migration, and mural-cell recruitment, the underlying transcriptional states of endothelial and perivascular cells remain unresolved. Integrating sustained perfusion with transcriptomic profiling will be an important next step toward understanding how flow-dependent changes in cell state give rise to the dynamic remodeling behaviors observed in angiogenic sprouts and how these processes contribute to vascular maturation.

Taken together, our results establish sustained laminar flow as a biologically meaningful regulator of angiogenic vascular organization in microphysiological systems. By combining tunable long-term perfusion with organotypic sprout architecture, endothelial polarity analysis, mural-cell recruitment, live imaging, and immune-cell interaction assays, this work provides a practical and experimentally accessible framework for studying how hemodynamic cues shape developing vascular networks on-chip. The modular nature of the workflow further suggests broad applicability to vascularized organ-on-chip systems, blood-brain barrier models, inflammatory assays, and future studies aimed at understanding how flow regulates vascular maturation and function in engineered tissues.

## Methods

### Cell culture

Pooled human umbilical vein endothelial cells from three donors, comprising two female and one male donor (HUVECs; PromoCell, C-12203) and human brain vascular pericytes (ScienCell, SC-1200) were cultured in tissue culture flasks coated with 0.2% gelatin (Merck, G1393). Cells were maintained in Endothelial Basal Medium-2 (EBM-2; Lonza, CC-3156) supplemented with the EGM-2 BulletKit (Lonza, CC-3162) at 37 °C in a humidified incubator with 5% CO₂. Cells were routinely passaged using standard procedures and experiments were performed using cells between passages 3 and 5.

### Vasculature-on-chip culture

Microfluidic chips (AIM Biotech, idenTx 3, DAX-1) were used to generate angiogenic vascular cultures. The central gel compartment was filled with a fibrin matrix prepared by mixing thrombin (Merck, T4393-100UN) and fibrinogen (Merck, F3879). Thrombin stock solution was reconstituted according to the manufacturer’s instructions and diluted in EBM-2 medium to a final concentration of 2 U/mL. The thrombin solution was then mixed 1:1 with fibrinogen solution (5 mg/mL) immediately before loading into the chip. Following loading, the gel was allowed to polymerize at room temperature for 8 min. After matrix polymerization, side channels were coated with 0.2% gelatin (Merck, G1393). HUVECs and pericytes were subsequently seeded into their respective channels. The HUVEC channel was maintained in EBM-2 supplemented with gentamicin-amphotericin, whereas the pericyte channel was maintained in EGM-2 supplemented with gentamicin-amphotericin and an additional 50 ng/mL VEGF-A (PeproTech, 100-20), thereby establishing a VEGF-A gradient across the gel compartment. Medium was exchanged every second day. Cultures were maintained for 14 days to allow the formation of lumenized angiogenic sprouts. Immediately before perfusion, the media in both side channels were replaced with the same complete EGM-2 medium without supplementary VEGF-A, thereby removing the exogenous VEGF-A gradient. The perfusion circuit was subsequently operated using the same complete EGM-2 medium.

### Laminar perfusion workflow

A modular perfusion system was assembled to provide tunable, unidirectional flow through microfluidic vasculature-on-chip cultures. Experiments were performed at a target flow rate of 30 μL/min for 24 h. The total circulating medium volume was approximately 5 mL per setup. The perfusion circuit consisted of a microfluidic chip, pulse damper (Bartels Mikrotechnik, BM-A-0018), unidirectional valve (Bartels Mikrotechnik, BM-A-0033), custom-made bubble trap, piezoelectric micropump (Bartels Mikrotechnik, mp6 and BP7), medium reservoir (Bartels Mikrotechnik, BM-A-0030), and inline flow sensor (Bartels Mikrotechnik, BM-S-0010) connected in series using 1.3 mm Tygon® tubing (Bartels Mikrotechnik, BM-A-0024). The custom bubble trap was assembled from a syringe barrel (Ibidi, 10967) and needle connector (Weller, KDS2312P). Flow was generated using the Bartels mp6 or BP7 piezoelectric micropump controlled through a Multiboard2 controller (Bartels Mikrotechnik) operating with Multiboard2 App v2.2 and firmware version 20230317 and equipped with the following pump drivers: mpDriver (Bartels Mikrotechnik, BM-E-0001), mp-Lowdriver (Bartels Mikrotechnik, BM-E-0004), mp-Highdriver4 (Bartels Mikrotechnik, BM-E-0003). Prior to connection to the microfluidic chips, the perfusion circuit was primed with the culture medium, the pump was primed by flushing 2 mL of media, and the circuit was subsequently operated at 1 mL/min for at least 2 h inside a humidified incubator maintained at 37 °C and 5% CO₂. This equilibration step facilitated medium degassing and removal of residual air bubbles from the system. Following equilibration, the microfluidic chips were connected to the perfusion circuit and returned to the incubator for experimental operation. To minimize leakage during long-term culture, the perfusion system was configured in a pull mode in which medium was drawn through the microfluidic device rather than pushed into it. The central gel-channel inlets were sealed and the pericyte-channel inlets were blocked, allowing medium to pass through the pericyte compartment while preventing reservoir depletion and unwanted leakage. Within each chip, the three endothelial channels were connected in series, enabling simultaneous perfusion of all technical replicates under identical flow conditions.

### Tracer bead perfusion imaging

To assess intraluminal perfusion, fluorescent 1 μm tracer beads (Invitrogen, F8816) were introduced into the perfusion circuit. Time-lapse imaging was performed using a MICA microscope (Leica Microsystems) equipped with a 10× HC PL FLUOTAR NA 0.32 objective and a 5 MP CMOS FluoSync^TM^. Images were acquired at 100 ms exposure, 1.6 Hz for 1 min. Representative frames were extracted from recorded image sequences to visualize bead movement through angiogenic sprouts.

### Immunofluorescence staining

For endpoint analysis, cultures were fixed with 4% paraformaldehyde (Merck, 158127) for 20 min at room temperature. Samples were subsequently blocked and permeabilized using blocking buffer containing 3% bovine serum albumin (Serva, 11930.03), 0.05% Triton X-100 (Sigma Aldrich, T8787-250ML), 0.01% sodium deoxycholate (Merck, D6750-25G), and 0.02% sodium azide (Merck, S2002-25G). Samples were incubated with primary antibodies against Ve-Cadherin, PDGFRβ, GM130, and Ki67 diluted in blocking buffer for 72 h at 4 °C. After washing, samples were incubated with fluorophore-conjugated secondary antibodies for 24 h at 4 °C. Nuclei were counterstained with DAPI (1:1000 in PBS, Life Technologies, D1306) for 20 min at room temperature. A complete list of antibodies and dilutions is provided in **Supplementary Table 1**.

### Polarity and orientation analysis

Endothelial cell orientation and Golgi-nuclear polarity were quantified using PolJam^36^. For each condition, three independent biological replicates were analyzed. The static dataset comprised 2,010 endothelial cells, whereas the flow dataset comprised 1,714 endothelial cells. For each microfluidic chip, two fields of view were acquired per endothelial channel, resulting in six analyzed images per chip. One field was positioned proximal to the flow inlet and the second proximal to the flow outlet. Maximum intensity projections were generated from acquired z-stacks and subjected to background subtraction in Fiji (ImageJ, National Institutes of Health). Cell segmentation was performed using μSAM^46^. Nuclear position was identified using DAPI staining, while Golgi position was determined from GM130 immunostaining. Cell orientation and Golgi-nuclear polarity vectors were subsequently calculated using PolJam according to the developer’s recommendations. Rose plots, polarity indices, and Rayleigh tests of non-uniformity were generated using the PolJam analysis pipeline.

### Pericyte recruitment quantification

Pericyte recruitment was quantified by determining the number of PDGFRβ-positive pericytes associated with endothelial sprouts and normalizing this value to vessel area. Images were acquired as 2 × 2 tile scans consisting of 43 optical sections per field of view. Two fields of view were collected from each endothelial channel, resulting in six analyzed images per microfluidic chip. One field was positioned proximal to the flow inlet and the second proximal to the flow outlet. Vessel area was quantified from maximum intensity projections generated from the acquired z-stacks. Images were background-subtracted in Fiji (ImageJ, National Institutes of Health) and endothelial structures were segmented using μSAM^46^. The resulting vessel masks were used to calculate total vessel area for normalization. Because of the complex morphology of pericytes and the discontinuous distribution of PDGFRβ signal, vessel-associated pericytes were quantified manually according to predefined criteria. A pericyte was considered vessel-associated only if: 1) the entire cell body was in direct contact with an endothelial structure, 2) the associated Ve-Cadherin-positive vessel segment was in focus, and 3) a corresponding DAPI-positive nucleus could be identified. Cells that were partially detached from the vessel or lacked an identifiable nucleus were excluded from analysis. A total of 18 images, per condition, obtained from 3 biological replicates were analyzed. The number of vessel-associated pericytes was normalized to the segmented vessel area. Each data point shown in the quantification corresponds to one analyzed image.

### Ki-67 quantification

Ki-67 staining was quantified from z-stack images acquired using the MICA imaging platform (Leica Microsystems) and a 10× objective. For each field of view, 4 regions of interest (ROIs) were analyzed, typically comprising two ROIs from the lower side and two ROIs from the upper side of the mother vessel. Maximum intensity projections were generated from the acquired z-stacks prior to analysis. Ki-67-positive nuclei were identified manually using a standardized look-up table (LUT) applied to the Ki67 channel. Only nuclei showing clear Ki-67 signal and a corresponding DAPI-positive nuclear signal were counted as Ki-67-positive. To determine the total nuclear count, the DAPI channel was background-subtracted and thresholded to improve nuclear segmentation. The thresholded DAPI images were then analyzed in Fiji (ImageJ, National Institutes of Health) using particle analysis to count total nuclei. The percentage of proliferating HUVECs was calculated as the number of Ki-67-positive nuclei divided by the total number of DAPI-positive nuclei within each ROI. Each data point represents one analyzed ROI.

### Live-cell imaging

Live-cell imaging experiments were performed using a MICA microscope (Leica Microsystems) equipped with environmental control for temperature, humidity, and CO₂ regulation. Microfluidic chips were maintained at 37 °C and 5% CO₂ throughout image acquisition. For imaging of angiogenic sprout dynamics, microfluidic chips were mounted directly onto the microscope stage and imaged using a 10× objective. Time-lapse sequences were acquired using the integrated modular contrast setting at 40 ms exposure, every 5 min for 12 h. Images were processed using Fiji (ImageJ) and representative image sequences were exported for figure preparation and video generation. Live-cell imaging experiments were used to visualize endothelial migration, pericyte behavior, lumen formation, tip-cell extension, and immune-cell interactions under flow conditions.

### PBMC isolation, stimulation, and perfusion

Peripheral blood mononuclear cells (PBMCs) were isolated from leukocyte reduction filters obtained from the Charité Blood Donation Center under institutional guidelines and ethical approval (Approval Number: EA4/099/20). PBMCs were isolated according to standard laboratory procedures and cultured overnight in complete medium. Prior to perfusion experiments, PBMCs were stimulated with staphylococcal enterotoxin B from Staphylococcus aureus (SEB; Merck KGaA, S4881) for 16 h. SEB was diluted in sterile distilled water to achieve a final stock concentration of 10 µg/mL and was used for PBMC stimulation at a concentration of 0.3 µg/mL. For inflammatory perfusion assays, vasculature-on-chip cultures were stimulated with tumor necrosis factor alpha (3.3 ng/mL TNFα; Gibco, PHC3015) for 24 h before PBMC (5 × 10^6^ cells/mL medium) introduction. TNFα was maintained in the circulating medium throughout the experiment. Following stimulation, PBMCs were introduced into the medium reservoir and perfused through the vascular network under continuous flow conditions. Cultures were maintained under flow for 18 h prior to imaging. Time-lapse imaging was subsequently performed during the final 4 h of perfusion using the MICA microscope. The mother vessel was imaged using a 20× objective with a 40 ms exposure at 1 Hz for 10 min. Angiogenic sprouts were imaged using a 10× objective with a 40 ms exposure at 1 Hz for 5 min.

### Flow calibration

Flow rates were monitored using an inline flow sensor (Bartels Mikrotechnik, BM-S-0010) integrated into the perfusion circuit. Flow was adjusted through mp6 or BP7 control software to achieve a target flow rate of 30 μL/min. Following equilibration and chip connection, flow stability was confirmed before initiation of biological experiments. Importantly, the perfusion workflow is readily tunable and is not restricted to a single operating condition. Depending on the selected pump driver, flow rates can be continuously adjusted from 5 to 1,000 μL/min using the mp-Lowdriver and up to 9,000 μL/min using the mp-Highdriver4. This wide operating range enables the generation of diverse hemodynamic conditions and allows the same platform to be adapted to different microfluidic devices, vessel geometries, and experimental applications requiring either physiological or supraphysiological flow regimes.

### Reynolds-number estimate

To estimate the bulk flow regime within the mother-vessel channel, the Reynolds number was calculated as Re = ρUDₕ/μ, where ρ and μ are the density and dynamic viscosity of the culture medium, U is the mean channel velocity, and Dₕ is the hydraulic diameter. For the rectangular channel, U was calculated as Q/(wh) and Dₕ as 2wh/(w + h). Using the nominal channel width (w = 0.50 mm), height (h = 0.25 mm), and imposed flow rate (Q = 30 µL/min), the mean velocity and hydraulic diameter were estimated as 4.0 mm/s and 0.333 mm, respectively. Assuming a medium density of 993 kg/m³ and a water-like dynamic viscosity of 0.69–1.0 mPa·s at 37 °C, the resulting Reynolds number was approximately 1.3–1.9, indicating strongly laminar bulk flow within the mother-vessel channel. This estimate characterizes the nominal mother-channel flow regime and does not resolve local velocities or wall shear stresses within the angiogenic sprouts.

### Statistics and reproducibility

Data are presented as mean ± standard deviation (s.d.) unless otherwise indicated. Statistical analyses were performed using GraphPad Prism version 10. The statistical tests used for individual datasets are indicated in the corresponding figure legends. Circular data obtained from polarity analyses were evaluated using Rayleigh tests of non-uniformity. A value of P < 0.05 was considered statistically significant. Unless otherwise stated, experiments were performed using three independent biological replicates. No statistical methods were used to predetermine sample size. Investigators were not blinded during data acquisition or analysis. Raw microscopy data were processed using Fiji (ImageJ, National Institutes of Health).

### Use of large language models (LLMs)

Large language models (LLMs), including ChatGPT (OpenAI), were used to assist with non-scientific aspects of manuscript preparation, including rephrasing existing text for clarity and readability.

## Data availability

Data supporting the findings of this study are available from the corresponding author upon reasonable request.

## Code availability

No code was generated to support this manuscript.

## Acknowledgements

The authors support diversity and inclusiveness in science and believe strongly in equality of human rights. The authors would like to thank Prof. Jens Kurrek (TU Berlin), Dr. Hannes Gonschior (MDC Innovation & Entrepreneurship), as well as members of AG Gerhardt (MDC) and AG Vajkoczy (Charité) for helpful suggestions and discussion. This work was supported by the Einstein Center 3R (EC3R, Einstein Foundation Berlin, EZ-2020-597-2) and by the Deutsche Forschungsgemeinschaft (DFG, German Research Foundation) through Collaborative Research Centre SFB1588, “Decoding and Targeting Mechanisms of Neuroblastoma Evolution” (project number 493872418).

## Contributions

Conceptualization: T.N. and H.G.; methodology: T.N., and J.S.; validation: T.N.; formal analysis: T.N.; investigation: T.N., K.M., J.Ku., J.Kr., K.K., L.J. and L.V.; resources: I.H., M.H. and H.G.; data curation: T.N. and J.Kr.; writing - original draft: T.N.; writing - reviewing and editing: T.N., K.K., M.H. and H.G.; visualization: T.N.; supervision: H.G.; funding acquisition: J.S. and H.G.

## Corresponding author

Correspondence to Prof. Dr. Holger Gerhardt and Dr. Tomasz Nawara

## Competing interests

T.N. received payment from Leica Microsystems under a consulting agreement that included the presentation of work related to this study and promotional content concerning a Leica microscopy system. The payment was used to cover conference-related expenses. Leica Microsystems had no role in the study design, data collection, data analysis, interpretation of the results, preparation of the manuscript, or decision to publish. The remaining authors declare no competing interests.

## Supplementary information

**Supplementary Movie 1.** Directional bead transport from mother vessel to pericyte compartment under laminar perfusion

**Supplementary Movie 2.** Directional bead transport from pericyte compartment to mother vessel under laminar perfusion

**Supplementary Movie 3.** Long-term live imaging of angiogenic vasculature-on-chip during sustained perfusion

**Supplementary Movie 4.** Endothelial migration against the direction of flow within the mother vessel

**Supplementary Movie 5.** Endothelial migration against the direction of flow within angiogenic sprouts

**Supplementary Movie 6.** Lumen formation and remodeling during sustained laminar perfusion

**Supplementary Movie 7.** Dynamic pericyte migration and projection extension toward angiogenic sprouts

**Supplementary Movie 8.** Limited PBMC-endothelial interactions in untreated mother vessels

**Supplementary Movie 9.** Increased PBMC crawling and endothelial association in inflamed mother vessels

**Supplementary Movie 10.** PBMC trafficking and crawling within angiogenic sprouts under inflammatory conditions

**Supplementary Table 1.** List of antibodies used in the study.

| Name | Host | Dilution | Supplier | Cat Number |
| --- | --- | --- | --- | --- |
| α-Ve-Cadherin | Goat | 1:250 | R&D Systems | AF938 |
| α-PDGFRβ | Rabbit | 1:100 | Cell Signaling | 3169 |
| α-GM130 | Mouse | 1:300 | BD BioSciences | 610822 |
| α-Ki67-eFluor™ 660 | Rat | 1:100 | eBioscience | 50-5698-82 |
| Alexa Fluor 488 αRabbit | Donkey | 1:400 | ThermoFisher | A21206 |
| Alexa Fluor 568 αMouse | Donkey | 1:400 | ThermoFisher | A10037 |
| Alexa Fluor 568 αGoat | Donkey | 1:400 | ThermoFisher | A11057 |
| Alexa Fluor 647 αGoat | Donkey | 1:400 | ThermoFisher | A21447 |

